# Assessment of the relationship between synaptic density and metabotropic glutamate receptors in early Alzheimer’s disease: a multi-tracer PET study

**DOI:** 10.1101/2024.09.21.614277

**Authors:** Elaheh Salardini, Ryan S. O’Dell, Em Tchorz, Nabeel B. Nabulsi, Yiyun Huang, Richard E. Carson, Christopher H. van Dyck, Adam P. Mecca

## Abstract

**Background:** The pathological effects of amyloid β oligomers (Aβo) may be mediated through the metabotropic glutamate receptor subtype 5 (mGluR5), leading to synaptic loss in Alzheimer’s disease (AD). Positron emission tomography (PET) studies of mGluR5 using [^18^F]FPEB indicate a reduction of receptor binding that is focused in the medial temporal lobe in AD. Synaptic loss due to AD measured through synaptic vesicle glycoprotein 2A (SV2A) quantification with [^11^C]UCB-J PET is also focused in the medial temporal lobe, but with clear widespread reductions is commonly AD-affected neocortical regions. In this study, we used [^18^F]FPEB and [^11^C]UCB-J PET to investigate the relationship between mGluR5 and synaptic density in early AD.

**Methods:** Fifteen amyloid positive participants with early AD and 12 amyloid negative, cognitively normal (CN) participants underwent PET scans with both [^18^F]FPEB to measure mGluR5 and [^11^C]UCB-J to measure synaptic density. Parametric *DVR* images using equilibrium methods were generated from dynamic. For [^18^F]FPEB PET, *DVR* was calculated using equilibrium methods and a cerebellum reference region. For [^11^C]UCB-J PET, *DVR* was calculated with a simplified reference tissue model – 2 and a whole cerebellum reference region.

**Result:** A strong positive correlation between mGluR5 and synaptic density was present in the hippocampus for participants with AD (*r* = 0.81, *p* < 0.001) and in the CN group (*r* = 0.74, *p* = 0.005). In the entorhinal cortex, there was a strong positive correlation between mGluR5 and synaptic in the AD group (*r* = 0.85, *p* <0.001), but a weaker non-significant correlation in the CN group (*r* = 0.36, *p* = 0.245). Exploratory analyses within and between other brain regions suggested significant positive correlations between mGluR5 in the medial temporal lobe and synaptic density in a broader set of commonly AD-affected regions.

**Conclusion:** Medial temporal loss of mGluR5 in AD is associated with synaptic loss in both medial temporal regions and more broadly in association cortical regions, indicating that mGluR5 mediated Aβo toxicity may lead to early synaptic loss more broadly in AD-affected networks. In CN individuals, an isolated strong association between lower mGluR5 and lower synaptic density may indicate non-AD related synaptic loss.

## Background

Alzheimer’s disease (AD) results in early and pronounced synaptic loss as a prominent pathological feature (1–4). Evidence supports a robust correlation between synaptic loss and level of cognitive impairment (5, 6), as determined by postmortem and brain biopsy studies, as well as synaptic positron emission tomography (PET) imaging (7–10). [^11^C]UCB-J was developed as a PET tracer for synaptic vesicle glycoprotein 2A (SV2A) in the past decade and has shown promising results in investigations of synaptic density in human studies, including studies of AD (11–13). [^11^C]UCB-J has a high in vivo affinity for SV2A, which resides within synaptic vesicles located at presynaptic terminals (14, 15). We have reported widespread reductions in synaptic density in the medial temporal lobe and in common AD-affected neocortical brain regions using [^11^C]UCB-J PET (7, 13, 16). This has been corroborated by multiple other groups (17–22). Glutamate is the primary excitatory neurotransmitter in the nervous system with ionotropic glutamate receptors being the main conduit for information transfer within the central nervous system (23). However pre- and postsynaptic metabotropic glutamate receptors (mGluRs) are commonly present and help with fine-tuning synaptic communication between neurons by regulating strength and timing of network activity (24). Metabotropic glutamate receptor subtype 5 (mGluR5) is a seven-transmembrane G protein-coupled receptor expressed in neurons and glial cells throughout the cortex and hippocampus that has a non-homogeneous distribution pattern (24–28). Based on mouse hippocampal neuron studies, mGluR5 have been considered primarily post-synaptic and involved in inducing long-term depression (LTD) at NMDAR synapses (26, 29, 30). However, more recent evidence indicates a heterogenous localization and function for mGluR5 with presynaptic, postsynaptic, and intracellular expression. Non-human primate studies indicate that mGluR5 is expressed in both presynaptic and postsynaptic terminals in the dorsolateral prefrontal cortex (31). Additionally, studies in rats demonstrate the existence of functional intracellular mGluR5 in hippocampus. Based on animal models of AD, it has been hypothesized that mGluR5 contributes to amyloid-β oligomer (Aβo) toxicity through various mechanisms. This includes facilitating the clustering of Aβo as an extracellular scaffold for mGluR5 – leading to Aβo-induced abnormal mGluR5 accumulation and subsequent increase in intracellular calcium levels and synaptic deterioration (32), as well as mGluR5 acting as a co-receptor with cellular prior protein (PRPc) and subsequent postsynaptic activation of the tyrosine kinase Fyn (33, 34). A The latter finding asserts mGluR5 as a link between Aβ and tau pathology where the activation of Fyn leads to downstream tau phosphorylation (35). Recognition of mGluR5 as a mediator of AD pathology has spurred research into its role as a therapeutic target in AD mouse models as well as in human clinical trials (36–42).

Several recent human PET imaging studies with mGluR5 specific radiotracers have made it possible to assess mGluR5 changes in individuals affected by clinical AD. Our previous work quantifying mGluR5 availability in AD with [^18^F]FPEB PET showed a significant reduction of hippocampal mGluR5 due to AD with non-significant, but numerically lower mGluR5 binding in association cortical regions (25). This finding was corroborated in studies by Wang *et al.* and Treyer *et al.* using [^18^F]PSS232 and [^11^C]-ABP699 PET respectively (43, 44).

As an extension of our previous work showing synaptic density and mGluR5 reductions in AD we performed analyses to investigate the spatial relationships between both biomarkers in a cohort of individuals who underwent both [^18^F]FPEB and [^11^C]UCB-J PET. Because the largest reductions of mGluR5 and synaptic density are found in the medial temporal lobe, in our primary analyses we focused on the hippocampus and entorhinal cortex. We then examined brain wide regional correlations between mGluR5 and synaptic density. We hypothesized that mGluR5 and synaptic density would be strongly correlated in participants with AD, not in CN participants.

## Methods

The study protocol was approved by the Yale University Human Investigation Committee and Radiation Safety Committee. All participants provided written informed consent prior to participating in the study.

### Study Participants

Participants between 55 and 85 years of age were evaluated with a screening diagnostic evaluation, as previously described (45). Participants with AD were required to either i) meet the diagnostic criteria for probable dementia based on National Institute on Aging-Alzheimer’s Association (NIA-AA) guidelines, have a Clinical Dementia Rating (CDR) score of 0.5 -1, and Mini-Mental Status Examination (MMSE) score of ≥16 or ii) meet the NIA-AA diagnostic criteria of amnestic mild cognitive impairment (aMCI), have a CDR score of 0.5, and an MMSE score of 24-30. Moreover, participants in the AD group were required to demonstrate impaired episodic memory, as evidenced by a Logical Memory (LM) II score of 1.5 standard deviations (SD) below an education-adjusted norm. CN participants were required to have a CDR score of 0, an MMSE score > 26, and a normal education adjusted LMII.

All participants underwent PET with [^11^C]Pittsburg Compound B ([^11^C]PiB) to assess for the presence of brain Aβ. [^11^C]PiB PET scans were required to be negative for Aβ in CN participants and positive in AD participants. Participants were considered Aβ+ if the [^11^C]PiB PET scan was positive based on visual interpretation of 2 expert readers and confirmed with quantitative read criteria of cerebral-to-cerebellar distribution volume ratio (*DVR*) of at least 1.40 in at least 1 AD-affected region of interest (ROI) (7, 46).

### Magnetic resonance imaging

Magnetic resonance imaging (MRI) was conducted using a 3T Trio (Siemens Medical Systems, Erlangen, Germany) equipped with a circularly polarized head coil. MRI acquisition consisted of a Sag 3D magnetization-prepared rapid gradient-echo (MPRAGE) sequence with the following parameters: 3.34-msec echo time, 2500-msec repetition time, 1100-msec inversion time, 7-degree flip angle, and 180 Hz/pixel bandwidth. The resulting images have dimensions of 256×256×176 with a pixel size of 0.98× 0.98×1.0mm. The MRI procedure was used to make sure that patients did not show evidence of infection, infarction, or other brain lesions. Moreover, it served to delineate brain anatomy, assess atrophy, and perform partial volume correction (PVC) of PET images. Version 6.0 of FreeSurfer (http://surfer.nmr.mhg.harvard.edu/) was used to reconstruct cortical regions and perform volumetric segmentation used to define ROIs in participant native space (47).

### Positron emission tomography methods

PET images were acquired on the High-Resolution Research Tomograph (Siemens Medical Solution, Knoxville, TN, USA, 207 slices, resolution < 3 mm full width half maximum) (48). Dynamic [^11^C]PiB scans were obtained over a period of 90 minutes after the bolus administration of a tracer dose of up to 555MBq (49). [^18^F]FPEB was used to quantify regional brain availability of mGluR5. Using the previously evaluated bolus plus constant infusion paradigm (Kbol = 190 min), dynamic [^18^F]FPEB scans were taken for 60 minutes, beginning at 60 minutes after the initial injection of up to 185 MBq of tracer. Lastly, [^11^C]UCB-J PET was used for evaluating synaptic density by acquiring dynamic scans up to 90 minutes after administration of a tracer bolus of up to 740 MBq (50).

Using the Motion-compensation OSEM List-mode Algorithm for Resolution-recovery (MOLAR), list-mode data was reconstructed with event-by-event motion correction based on Vicra optical detector (NDI Systems, Waterloo, Canada) (51, 52). Software motion correction was applied to the dynamic PET images using a mutual-information algorithm (FSL-FLIRT, FSL 3.2; Analysis Group, FMRIB, Oxford, UK) to perform frame-by-frame registration to a summed image (0 - 10 min for [^11^C]UCB-J and 60 - 70 min for [^18^F]FPEB. A summed motion corrected PET image was used to create a registration between the MRI and PET scans for each participant. This PET to MRI registration was used to apply participant specific ROIs to parametric PET images.

For [^11^C] PiB, parametric images of binding potential (*BP*_ND_), the ratio at equilibrium of specifically bound radioligand to that of nondisplaceable radioligand in tissue, were generated using simplified reference tissue model–2 (SRTM2) using whole cerebellum as reference region. In order to account for potential partial volume effects, we performed partial volume correction of dynamic series for [^18^F]FPEB and [^11^C]UCB-J using the iterative Yang method (53, 54). Kinetic modeling was performed both with and without PVC of dynamic PET series. For [^18^F]FPEB image analysis, parametric images of *DVR* were generated with equilibrium methods using data collected from 90 to 120 minutes post bolus injection and a whole cerebellum reference region, as previously described (25). Lastly, for [^11^C]UCB-J, SRTM2 was applied to generate parametric *BP*_ND_ images PET frames from 0 to 60 minutes post injection and a whole cerebellum reference region.(55) For [^11^C]UCB-J, *BP*_ND_ was converted to *DVR* using the formula *DVR* = *BP*_ND_ + 1.(16, 49)

Reported values for each ROI are bilateral regions except where specified as left or right hemisphere. ROIs used for the composite of AD-affected brain regions are defined in Supplementary Table 1. ROIs used for the medial temporal composite included bilateral hippocampus, entorhinal cortex, parahippocampal cortex, and amygdala.

### Statistical analysis

Statistical analyses were performed using MATLAB R2018b (Mathworks, Natick, MA, USA) and SPSS 28 (IBM Corp, Armonk, NY). Between group comparisons were performed using χ^2^ tests for categorical variables, independent two-tailed t tests for continuous variables, as well as Mann-Whitney U tests for CDR global and CDR sum of boxes scores. Separate univariate regression analyses were used to evaluate the relationship between mGluR5 and synaptic density with the primary analysis focused on hippocampus and entorhinal cortex. Pearson’s correlation coefficients (*r*) and associated two-tailed *p* values were calculated to assess the strength of linear correlation between mGluR5 and synaptic density in each ROI, as well as a medial temporal composite region. Fisher *r*-to-*z* transformation was used to compare the strength of correlation of mGluR5 and synaptic density between AD and CN groups. Significant *p* value was defined as *<* 0.05. Analyses were performed both without and with PVC of PET data. Analyses including all brain regions did not include correction for multiple comparisons due to the exploratory nature of these investigations.

## Results

### Participants characteristics

The study sample consisted of 15 amyloid positive participants with AD and 12 amyloid negative participants with normal cognition. Demographic characteristics, cognitive assessment results, *APOE* genotype, and PET *DVR* measures for each group are shown in Table 1. Diagnostic groups were well balanced for age, sex, and education. Additionally, all participants with AD demonstrated typical clinical characteristics of aMCI or mild dementia, with significant deficits indicated by MMSE (24.1 ± 3.9), CDR global score (0.7 ± 0.2), and CDR sum-of-boxes score (4.0 ± 2.2) in comparison to participants with normal cognition (Table 1). *APOE* genotypes reflected expected patterns with higher copy numbers of the ε4 in the AD participant group. As expected from our previous studies, synaptic density ([^11^C]UCB-J PET *DVR*) was lower in both the hippocampus and a composite of common AD-affected brain regions in participants with AD compared to the CN group. In this slightly smaller sample than our previous study with [^18^F]FPEB PET (25), hippocampal mGluR5 density was lower, but not significantly different, in participants with AD compared to the CN group. In line with our previous observation, mGluR5 density in the composite of AD-affected regions was not lower in participants with AD compared to the CN group.

**Table 1.**
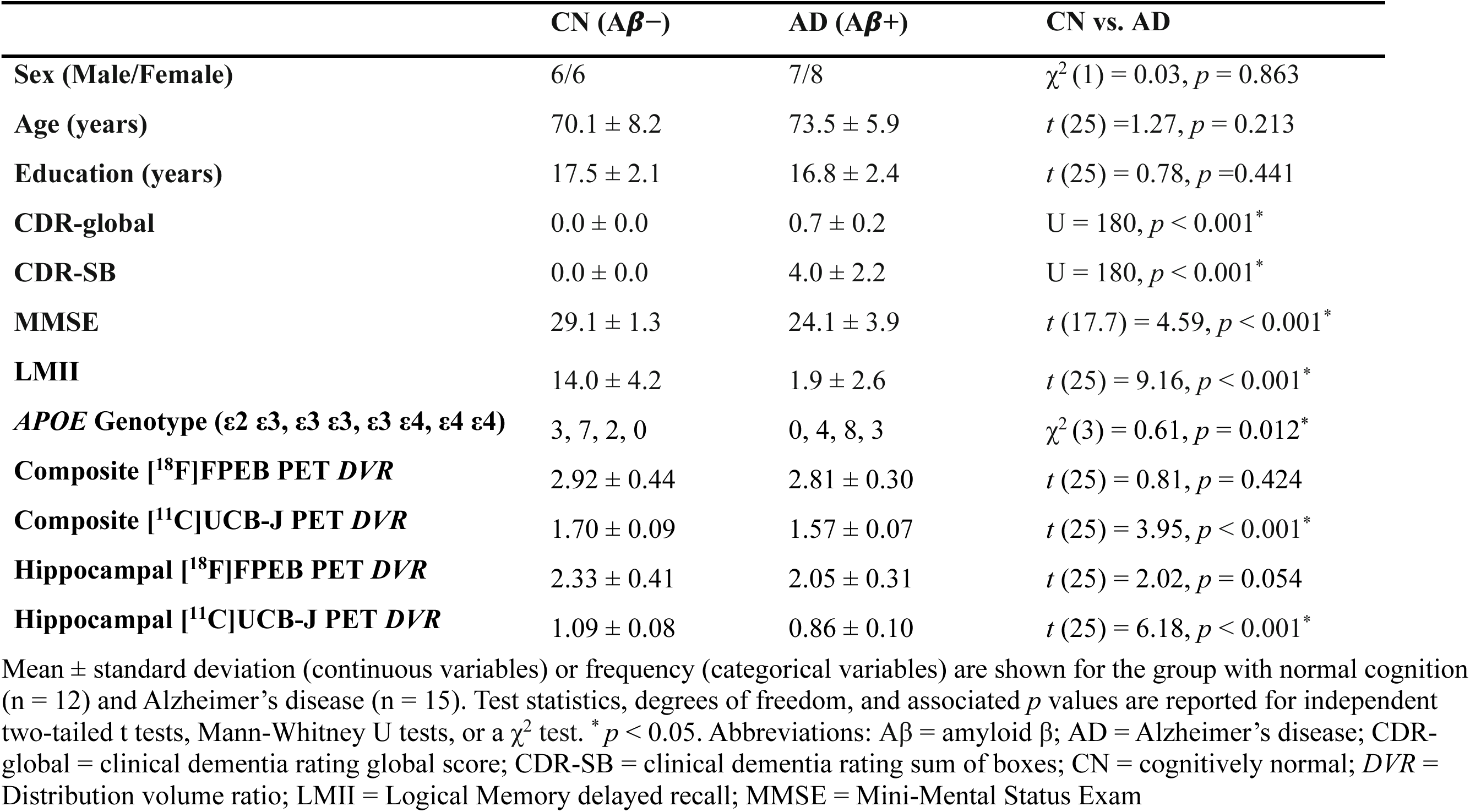
Participant characteristics.

### Correlations between mGluR5 and synaptic density in hippocampus and entorhinal cortex

Our primary analyses used univariate linear regression to assess the relationship between mGluR5 and synaptic density in the hippocampus and entorhinal cortex, regions known to be involved in early AD pathogenesis and with significant AD related reductions of synaptic density and mGluR5 density based on our previous studies. A strong, significant positive correlation was demonstrated between hippocampal mGluR5 and synaptic density in participants with AD (*r* = 0.81, *p* < 0.001) and a slightly weaker, significant positive correlation in the CN group (*r* = 0.74, *p* = 0.005, Figure 1A). Significant correlations of similar strength were also present in the hippocampus with PVC of the PET data (*r* = 0.82, *p* <0.001 for AD and *r* = 0.73, *p* = 0.007 for CN). A Fisher *r*-to-*z* transformation indicated no statistically significant difference in the strength of correlations in the hippocampus between the two groups without PVC (*z* = 0.35, *p* = 0.704) and with PVC (*z =* 0.50, *p* = 0.617). In the entorhinal cortex, a strong, significant positive correlation was demonstrated between mGluR5 and synaptic density in participants with AD (*r* = 0.85, *p* <0.001), but no significant correlation was found in the CN group (*r* = 0.36, *p* = 0.245, Figure 1B). Correlations of similar strength were also present in the entorhinal cortex with PVC of the PET data (*r* = 0.83 with *p* < 0.001 for AD, *r* = 0.40, *p* = 0.196 for CN). Although group differences in correlation strength were similar in magnitude, the correlation between mGluR5 and synaptic density in the entorhinal cortex was significantly stronger in participants with AD compared to the CN group without PVC (*z* =1.99, *p* = 0.046), but not with PVC (*z* = 1.76, *p* = 0.078).

**Figure 1.**
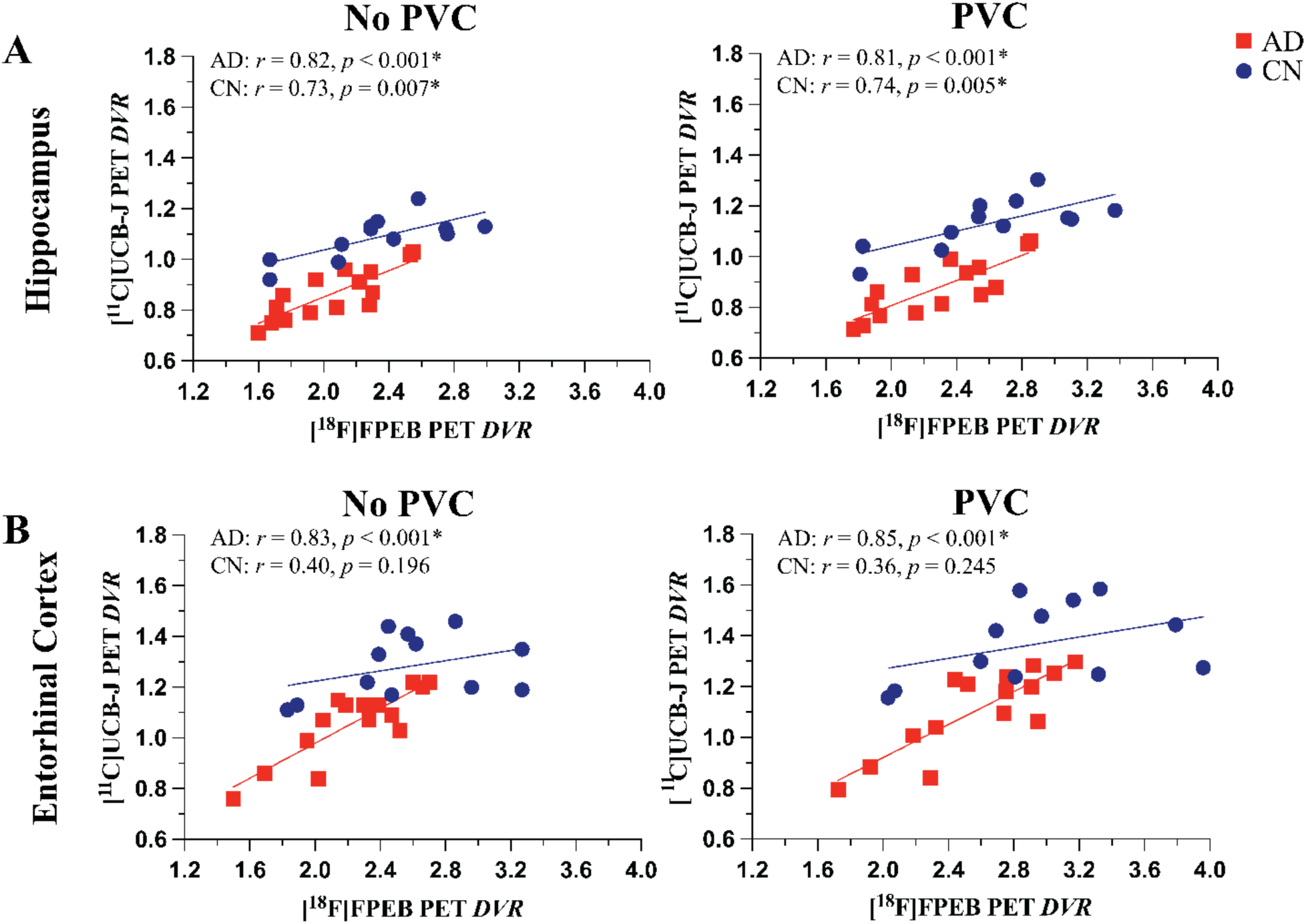
Correlations between mGluR5 and synaptic density in the hippocampus and entorhinal cortex. [^18^F]FPEB (mGluR5) and [^11^C]UCB-J (synaptic density) *DVR*s are plotted for participants with CN (blue, n = 12) and AD (red, n = 15). Univariate linear regression line of best fit, Pearson’s correlation coefficients (*r*) and the associated *p* values are shown for each group for **(A)** hippocampus and **(B)** entorhinal cortex. \**p* < 0.05. Abbreviations: AD = Alzheimer’s disease; CN = cognitively normal; *DVR =* Distribution volume ratio; mGluR5 = metabotropic glutamate receptor subtype 5; PVC = partial volume correction

### Correlations between mGluR5 and synaptic density in other medial temporal regions

To better understand the pattern of correlations in brain areas affected in early AD, we next focused on the medial temporal lobe in analyses of a composite medial temporal region, as well as the individual regions used to construct the composite (Supplementary Figure 1). There was a strong, positive correlation between mGluR5 and synaptic density in the medial temporal lobe of the AD group (*r* = 0.84, *p* < 0.001), and no significant correlation in the CN group (*r* = 0.56, *p* = 0.055). In addition to the relationships described in the primary analyses for hippocampus and entorhinal cortex, mGluR5 and synaptic density had a strong, positive correlation in the amygdala for the AD group (*r* = 0.84, *p* < 0.001), and a weaker non-significant correlation in the CN group (*r* = 0.56, *p* = 0.057). In the parahippocampal cortex, there was a strong, positive correlation in the AD group (*r* = 0.85, *p* < 0.001) and a weaker, non-significant correlation in the CN group (*r* = 0.42, *p* = 0.170). A very similar pattern of correlation and significance existed with application of PVC to the PET data, as well as after adjustment for multiple comparisons using false discovery rate (Supplementary Figure 1). All statistically significant correlations in the figure remained statistically significant after correction for multiple comparisons using false discovery rate (FDR) method except for amygdala in data with PVC in participants with normal cognition.

### Correlations between mGluR5 and synaptic density in all brain regions

We performed exploratory analyses in all brain regions to have a better understanding of the whole brain pattern of correlations between mGluR5 and synaptic density. Stronger significant correlations between mGluR5 and synaptic density were observed more broadly in participants with AD compared to the CN group both without and with PVC (Figure 2, Table 2, and Supplementary Table 2). Without PVC, regions with significant correlations in the AD group included bilateral temporal poles, entorhinal cortices, hippocampi, parahippocampal cortices, amygdalae, fusiform gyri, inferior/middle/superior temporal gyri, banks of the superior temporal sulci, insular cortices, medial orbitofrontal cortices, rostral anterior cingulate gyri, as well as left caudate, right pars opercularis, right transverse temporal gyrus, right supramarginal gyrus, right isthmus of the cingulate, right inferior parietal cortex, and right lingual gyrus. In the CN group, significant correlations existed only in the bilateral hippocampi, bilateral caudate, left pallidum, right transverse temporal cortex, left insular cortex, and left thalamus. When using Fisher *r*-to-*z* transformation to assess the difference in correlation strength between AD and CN groups, the bilateral temporal poles, left banks of superior temporal sulcus, and right entorhinal cortex had significantly stronger positive correlations in participants with AD as compared to the CN group. Similar relationships were seen with and without PVC of PET data (Table 2 and Supplementary Table 2).

**Figure 2.**
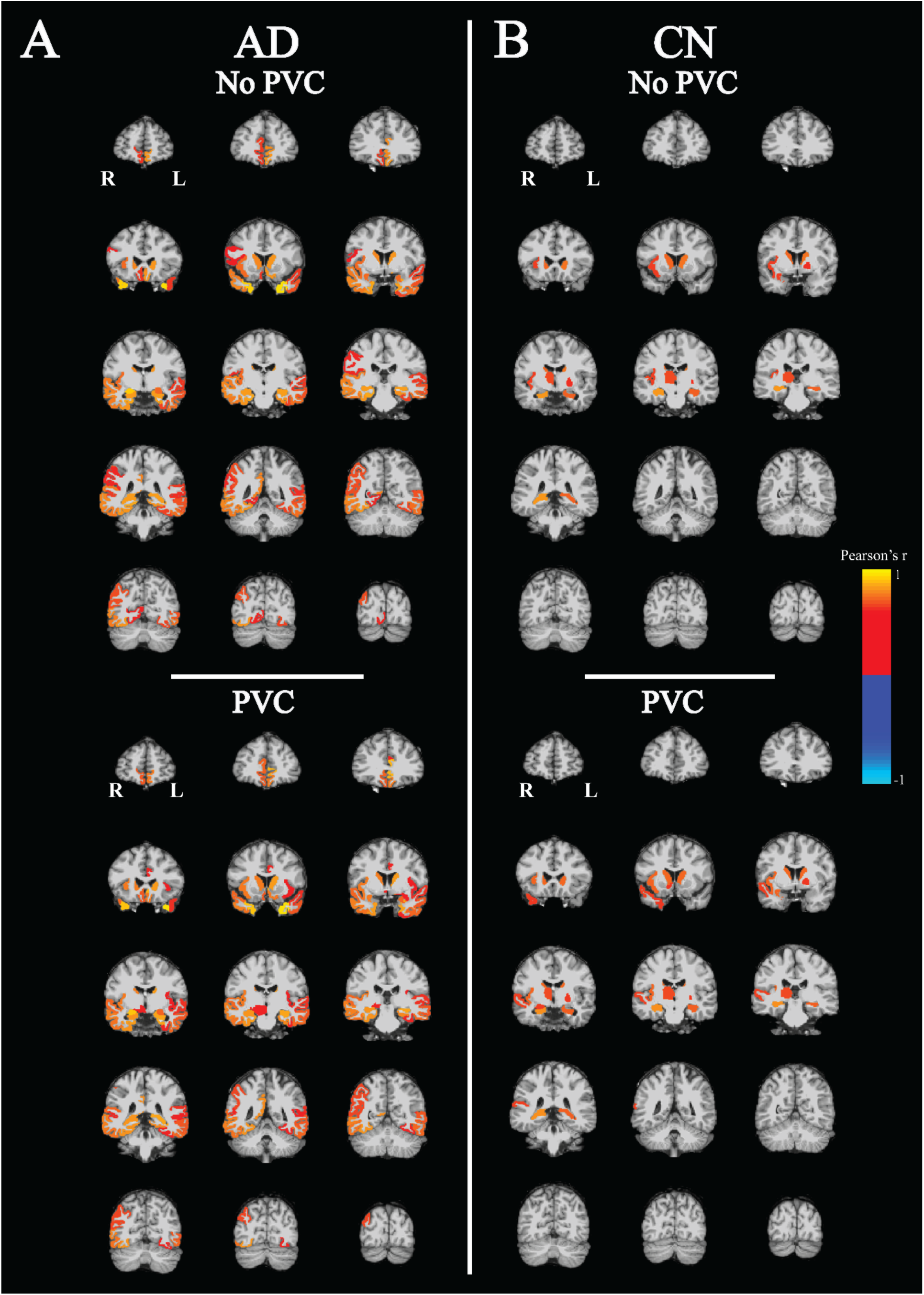
Correlation maps of mGluR5 and synaptic density in all regions. Pearson’s correlation coefficients (*r*) and associated *p* values were calculated between [^18^F]FPEB (mGluR5) and [^11^C]UCB-J (synaptic density) PET *DVRs* in all regions in participants with **(A)** Alzheimer’s Disease (n = 15) and **(B)** normal cognition (n = 12). All voxels in each region were colored uniformly for regions that had an uncorrected *p* < 0.05 and displayed as an overlay on the MNI template T1 MRI. Abbreviations: AD = Alzheimer’s disease; CN = cognitively normal; *DVR =* Distribution volume ratio; mGluR5 = metabotropic glutamate receptor subtype 5; PVC = partial volume correction

**Table 2.** Regional correlations between mGluR5 and synaptic density (no PVC)

| Region | Left hemisphere |  |  |  |  |  | Right hemisphere |  |  |  |  |  |
| --- | --- | --- | --- | --- | --- | --- | --- | --- | --- | --- | --- | --- |
| | AD | | CN | | Fisher's $r$ to $z$ | | AD | | CN | | Fisher's $r$ to $z$ | |
| | $r$ | $p$ | $r$ | $p$ | $z$ | $P$ | $r$ | $p$ | $r$ | $p$ | $z$ | $P$ |
| Frontal pole | 0.30 | 0.283 | 0.09 | 0.771 | 0.48 | 0.631 | 0.40 | 0.143 | -0.37 | 0.229 | 1.85 | 0.064 |
| Superior frontal | -0.03 | 0.905 | 0.05 | 0.881 | 0.19 | 0.852 | 0.02 | 0.927 | -0.02 | 0.939 | 0.11 | 0.909 |
| Rostral middle frontal | 0.13 | 0.650 | -0.08 | 0.804 | 0.47 | 0.636 | 0.31 | 0.255 | 0.15 | 0.635 | 0.39 | 0.699 |
| Caudal middle frontal | 0.10 | 0.715 | -0.17 | 0.590 | 0.63 | 0.528 | 0.39 | 0.152 | 0.08 | 0.813 | 0.76 | 0.449 |
| Pars orbitalis | 0.23 | 0.406 | 0.49 | 0.108 | 0.67 | 0.503 | -0.10 | 0.720 | 0.19 | 0.554 | 0.66 | 0.506 |
| Pars opercularis | 0.21 | 0.453 | 0.36 | 0.244 | 0.38 | 0.702 | 0.52 | 0.048* | 0.44 | 0.149 | 0.22 | 0.827 |
| Pars triangularis | 0.24 | 0.397 | 0.33 | 0.287 | 0.24 | 0.807 | 0.37 | 0.171 | 0.47 | 0.126 | 0.26 | 0.796 |
| Lateral orbitofrontal | 0.40 | 0.136 | 0.12 | 0.714 | 0.70 | 0.483 | 0.29 | 0.290 | 0.46 | 0.136 | 0.43 | 0.663 |
| Medial orbitofrontal | 0.79 | <0.001* | 0.44 | 0.147 | 1.35 | 0.178 | 0.59 | 0.020* | 0.49 | 0.101 | 0.31 | 0.759 |
| Temporal pole | 0.93 | <0.001* | 0.36 | 0.247 | 2.99 | 0.003* | 0.91 | <0.001* | 0.53 | 0.076 | 2.12 | 0.034* |
| Entorhinal | 0.81 | <0.001* | 0.46 | 0.127 | 1.42 | 0.155 | 0.83 | <0.001** | 0.18 | 0.575 | 2.27 | 0.023* |
| Parahippocampal | 0.78 | <0.001* | 0.46 | 0.135 | 1.27 | 0.205 | 0.83 | <0.001* | 0.36 | 0.245 | 1.82 | 0.068 |
| Hippocampus | 0.81 | <0.001* | 0.68 | 0.014 | 0.64 | 0.523 | 0.80 | <0.001* | 0.80 | 0.002* | 0.05 | 0.960 |
| Amygdala | 0.70 | 0.003* | 0.49 | 0.108 | 0.77 | 0.440 | 0.90 | <0.001* | 0.56 | 0.056 | 1.87 | 0.061 |
| Inferior temporal | 0.71 | 0.003* | 0.38 | 0.225 | 1.13 | 0.259 | 0.74 | 0.002* | 0.34 | 0.278 | 1.35 | 0.177 |
| Fusiform | 0.62 | 0.014* | 0.26 | 0.420 | 1.04 | 0.298 | 0.79 | <0.001* | 0.29 | 0.355 | 1.75 | 0.081 |
| Middle temporal | 0.65 | 0.008* | 0.23 | 0.464 | 1.23 | 0.220 | 0.76 | <0.001* | 0.49 | 0.107 | 1.07 | 0.286 |
| Bankssts | 0.57 | 0.026* | -0.43 | 0.162 | 2.51 | 0.012* | 0.60 | 0.017* | 0.46 | 0.131 | 0.45 | 0.652 |
| Superior temporal | 0.59 | 0.019* | 0.25 | 0.435 | 0.97 | 0.329 | 0.74 | 0.002* | 0.57 | 0.050 | 0.67 | 0.504 |
| Transverse temporal | 0.37 | 0.175 | 0.18 | 0.572 | 0.46 | 0.643 | 0.75 | 0.001* | 0.61 | 0.033* | 0.60 | 0.545 |
| Supramarginal | 0.50 | 0.058 | 0.16 | 0.626 | 0.88 | 0.377 | 0.55 | 0.032* | 0.31 | 0.318 | 0.68 | 0.497 |
| Insula | 0.46 | 0.087 | 0.21 | 0.511 | 0.63 | 0.526 | 0.70 | 0.003* | 0.61 | 0.035* | 0.38 | 0.704 |
| Rostral anterior cingulate | 0.77 | <0.001* | 0.47 | 0.118 | 1.13 | 0.258 | 0.61 | 0.016* | 0.27 | 0.387 | 0.96 | 0.336 |
| Caudal anterior cingulate | 0.47 | 0.079 | 0.19 | 0.557 | 0.72 | 0.473 | 0.37 | 0.176 | 0.03 | 0.915 | 0.80 | 0.424 |
| Posterior cingulate | 0.34 | 0.219 | -0.40 | 0.198 | 1.75 | 0.079 | 0.38 | 0.158 | -0.24 | 0.460 | 1.46 | 0.144 |
| Isthmus cingulate | 0.45 | 0.093 | 0.42 | 0.175 | 0.08 | 0.934 | 0.75 | 0.001* | 0.24 | 0.453 | 1.68 | 0.092 |
| Precuneus | 0.27 | 0.335 | -0.24 | 0.458 | 1.17 | 0.242 | 0.44 | 0.098 | -0.02 | 0.949 | 1.12 | 0.260 |
| Paracentral | 0.07 | 0.811 | 0.002 | 0.993 | 0.15 | 0.883 | 0.28 | 0.315 | -0.14 | 0.656 | 0.98 | 0.329 |
| Postcentral | 0.009 | 0.973 | 0.24 | 0.450 | 0.54 | 0.592 | 0.25 | 0.360 | 0.13 | 0.683 | 0.30 | 0.772 |
| Precentral | 0.06 | 0.837 | -0.02 | 0.952 | 0.18 | 0.860 | 0.13 | 0.648 | 0.03 | 0.924 | 0.22 | 0.824 |
| Superior parietal | 0.20 | 0.468 | -0.27 | 0.397 | 1.09 | 0.275 | 0.14 | 0.605 | -0.10 | 0.763 | 0.55 | 0.580 |
| Inferior parietal | 0.27 | 0.325 | -0.08 | 0.807 | 0.81 | 0.416 | 0.64 | 0.010* | 0.15 | 0.640 | 1.38 | 0.167 |
| Lateral occipital | 0.15 | 0.600 | -0.29 | 0.351 | 1.03 | 0.304 | 0.35 | 0.204 | -0.16 | 0.628 | 1.18 | 0.238 |
| Cuneus | 0.21 | 0.440 | -0.36 | 0.246 | 1.36 | 0.174 | 0.34 | 0.208 | -0.03 | 0.926 | 0.88 | 0.376 |
| Pericalcarine | 0.20 | 0.477 | 0.38 | 0.225 | 0.44 | 0.656 | 0.44 | 0.102 | 0.21 | 0.512 | 0.58 | 0.559 |
| Lingual | 0.28 | 0.302 | -0.02 | 0.944 | 0.72 | 0.472 | 0.54 | 0.037* | 0.23 | 0.477 | 0.85 | 0.395 |
| Thalamus | 0.26 | 0.344 | 0.25 | 0.431 | 0.03 | 0.978 | 0.35 | 0.203 | 0.63 | 0.026* | 0.87 | 0.381 |
| Caudate | 0.79 | <0.001* | 0.72 | 0.008 | 0.35 | 0.722 | 0.76 | <0.001* | 0.70 | 0.012* | 0.33 | 0.738 |
| Putamen | 0.24 | 0.397 | 0.21 | 0.504 | 0.05 | 0.958 | 0.12 | 0.656 | 0.18 | 0.563 | 0.14 | 0.888 |
| Pallidum | 0.18 | 0.526 | 0.58 | 0.049 | 1.08 | 0.279 | 0.08 | 0.765 | 0.37 | 0.232 | 0.70 | 0.485 |
| Accumbens area | 0.33 | 0.230 | 0.07 | 0.820 | 0.61 | 0.542 | 0.41 | 0.126 | 0.57 | 0.052 | 0.48 | 0.632 |
| Ventral Diencephalon | 0.24 | 0.395 | 0.009 | 0.976 | 0.52 | 0.599 | 0.50 | 0.054 | 0.09 | 0.768 | 1.05 | 0.295 |

Pearson's $r$ and associated $p$ value is reported for the correlation between [ $^{18}\text{F}$ ]FPEB (mGluR5) and [ $^{11}\text{C}$ ]UCB-J PET (synaptic density) $DVR$ in each brain region. Fisher $r$ -to- $z$ transformation was used to compare correlation coefficients of CN and AD groups. The data were from 12 CN participants 15 participants with AD. \* $p < 0.05$ .
Abbreviations: AD = Alzheimer's disease; CN = cognitively normal; *DVR* = Distribution volume ratio; PVC = Partial volume correction

To explore the relationships of mGluR5 and synaptic density between different brain regions, we constructed a matrix of inter-tracer correlations for all region pairs in each diagnostic group. A review of these matrices with an overlaid heatmap of the correlation strength indicates strong correlations between synaptic density in the medial orbitofrontal and temporal lobes with mGluR5 in widespread brain regions in participants with AD (Figure 3). In the CN group, significant moderate to strong correlations were more isolated between hippocampal synaptic density and mGluR5 in widespread brain regions (Figure 4). Similar relationships existed with PVC of the PET images (Supplementary Figures 2 and 3).

**Figure 3.**
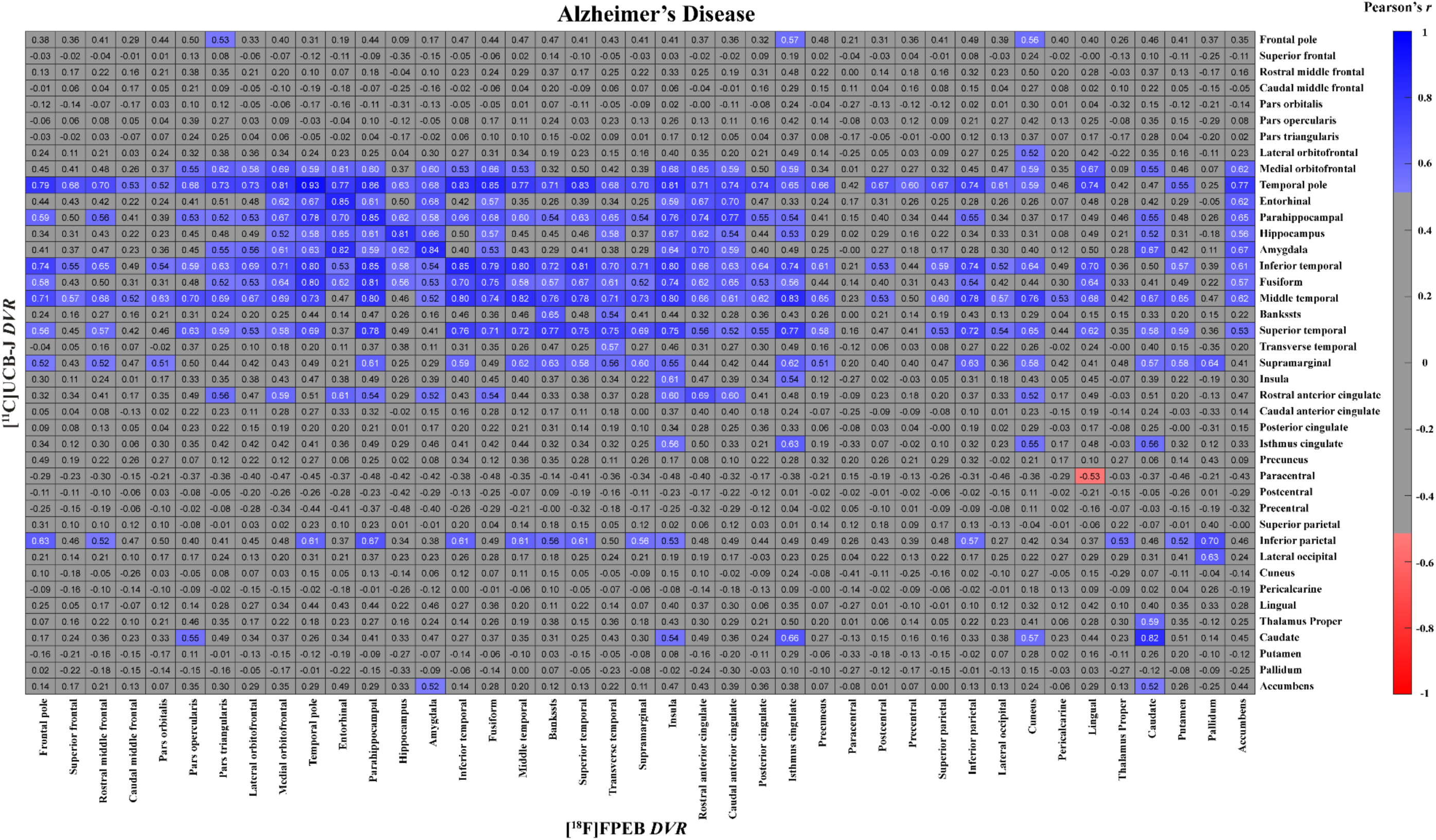
Correlation matrix for mGluR5 and synaptic density (no PVC) of all regions in participants with AD. The matrix displays Pearson’s correlation coefficients (*r*) between [^18^F]FPEB (mGluR5) and [^11^C]UCB-J (synaptic density) PET *DVRs* for all possible combinations of regions. Data are from 15 participants with Alzheimer’s Disease. The heat map shows the *r* for all combinations that had an uncorrected *p* < 0.05. Abbreviations: AD = Alzheimer’s disease; CN = cognitively normal; *DVR* = Distribution volume ratios; mGluR5 = metabotropic glutamate receptor subtype 5; PVC = partial volume corrected

**Figure 4.**
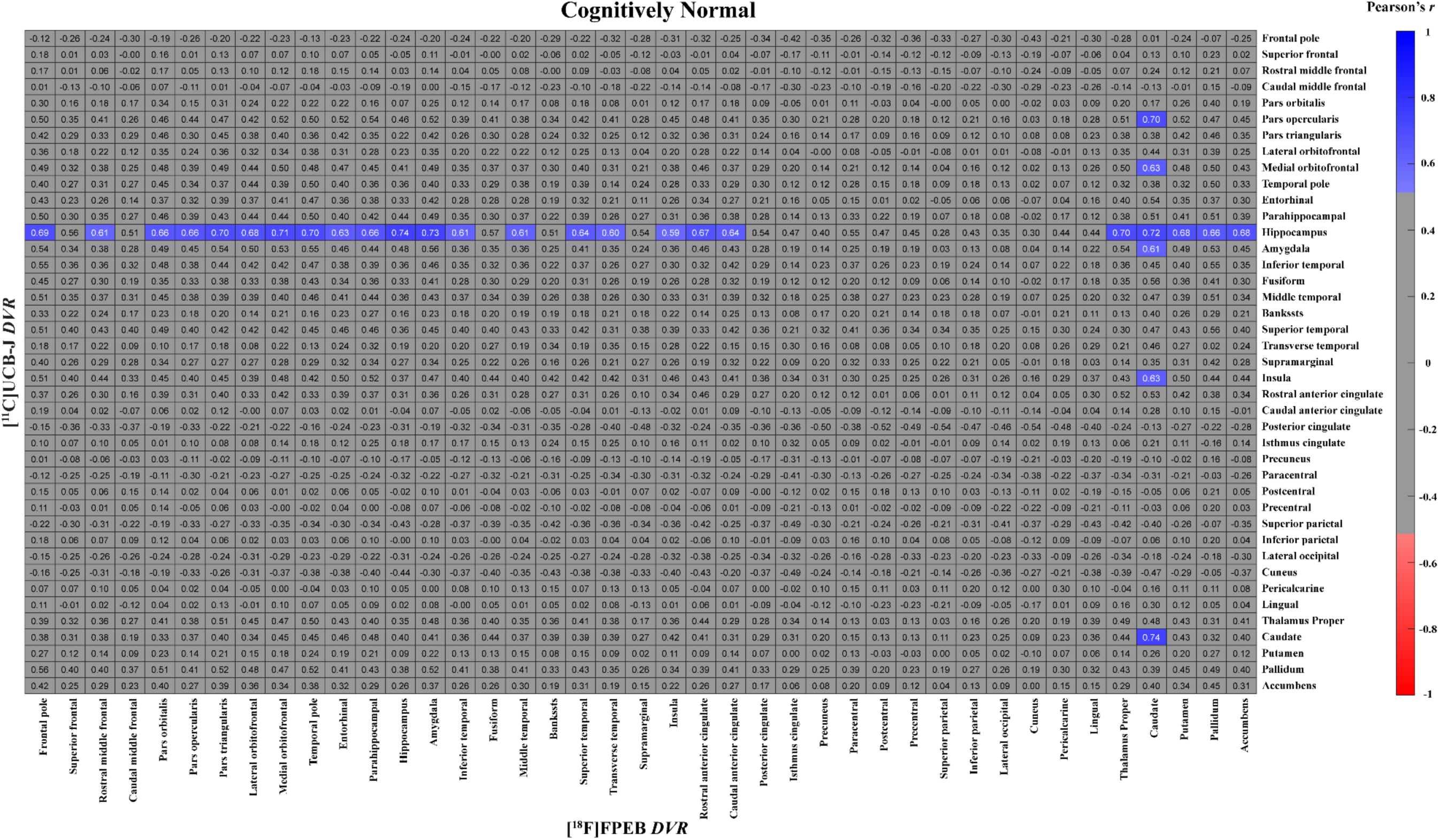
Correlation matrix for mGluR5 and synaptic density (no PVC) of all regions in CN participants. The matrix displays Pearson’s correlation coefficients (*r*) between [^18^F]FPEB (mGluR5) and [^11^C]UCB-J (synaptic density) PET *DVRs* for all possible combinations of regions. Data are from 12 cognitively normal participants. The heat map shows the *r* for all combinations that had an uncorrected *p* < 0.05. Abbreviations: AD = Alzheimer’s disease; CN = cognitively normal; *DVR* = Distribution volume ratios; mGluR5 = metabotropic glutamate receptor subtype 5; PVC = partial volume corrected

## Discussion

In this study we investigated the relationship between mGluR5 availability measured with [^18^F]FPEB PET and synaptic density measured with [^11^C]UCB-J binding to SV2A in early AD compared to individuals with normal cognition with an initial focus on medial temporal brain regions, followed by region-based whole brain analyses. We found strong correlations between mGluR5 binding and synaptic density in the hippocampus and entorhinal cortex in individuals with AD. This differed from the CN group where a strong positive correlation between mGluR5 availability and synaptic density was present in the hippocampus, but not the entorhinal cortex. In the whole brain region-based analyses, widespread significant positive correlations between mGluR5 binding and synaptic density were found in the group with AD.

Our previous work using [^18^F]FPEB and [^11^C]UCB-J PET showed that both mGluR5 binding and synaptic density are significantly lower in the medial temporal lobe of individuals with AD with largest effect sizes in the hippocampus (13). While the AD-related reduction in mGluR5 was significant only in the hippocampus the magnitude of mGluR5 was significantly lower in many commonly AD-affected association cortical regions (25). The synaptic density reduction due to AD was observed was of larger magnitude and more widespread in neocortical brain regions (7). Our results indicate that mGluR5 and synaptic density are highly correlated within a group of participants with early AD. Considering that synaptic density is highly correlated with cognitive performance in a larger sample of participants with AD (45),it is possible that loss of mGluR5 and SV2A are markers of disease progression that are highly related due to their locations at the synapse. Interestingly, mGluR5 binding and synaptic density were strongly correlated in the hippocampus, but not the entorhinal cortex in the CN group. This hippocampal correlation in the CN group was similar in magnitude and not significantly different in comparison to the group with AD. The meaning of this correlation in CN participants is not clear, but there may be non-AD-related reductions in mGluR5 and synaptic density in this group of older adults that are correlated, but not causing clinical symptoms.

We also investigated the within-region relationships between mGluR5 and synaptic density in all individual brain ROIs. We found that mGluR5 binding and synaptic density were significantly correlated with a widespread spatial extent in the AD group, but that intraregional correlations where more isolated in the CN group. In addition to the possibility that some of these intraregional relationships may be driven by non-AD disease processes – such as those in the hippocampus – it is also possible that age-related neurodegeneration could contribute in some regions. Of particular interest, we found the strongest correlation between mGluR5 binding and synaptic density in the CN group exists in the caudate. In our work and the work of others, the caudate has the strongest correlation between age and synaptic density, suggesting that this may be a site of age-related synaptic loss (56–58). We speculated that this association may be present because the caudate is the site of nerve terminals for multiple major tracts that undergo substantial age-related neurodegeneration (56). Similarly, mGluR5 binding and age are most strongly correlated in the caudate, although this age-related reduction in mGluR5 binding may be largely mediated by brain volume loss (59).

There is one other study investigating the relationship between mGluR5 binding measured with [^18^F]PSS232 PET and synaptic density measured with [^18^F]SynVesT-1 PET in a cohort of 20 participants (10 CN and 10 AD). In this study by Wang *et al*., they reported significant correlations between mGluR5 binding and synaptic density within and between many typically AD-affected regions and also performed a more speculative analysis that suggested mGluR5 binding in the medial temporal lobe may mediate the association between global amyloid and synaptic density in that region (60). The results of Wang *et al*. are not that surprising since their analysis was performed in the entire cohort of CN and AD participants and therefore, likely driven by the large differences in these groups due to the presence or absence of AD pathogenesis. A strength of our paper that builds on previous findings is separate analyses in CN and AD groups that may help distinguish AD from non-AD related associations between mGluR5 binding and synaptic density.

### Limitations

Our study has a few limitations. The diagnosis and stage of AD was determined with clinical criteria and amyloid PET positivity with no assessment of brain tau accumulation that may have provided a better understanding of AD pathological stage. Moreover, the relatively small sample size limits our ability to detect subtle relationships when signal-to-noise ratios may be low. Future studies with larger sample sizes could confirm the absence of correlations, and also allow investigations into the relationship between mGluR5 and synaptic density with cognition. Additionally, our study is cross-sectional which limits the ability to determine causal relationships. Longitudinal assessments with both radiotracers starting at preclinical AD stages would allow validation of findings and a more thorough investigation of the temporal and spatial changes of mgluR5 and synaptic density due to AD progression.

### Conclusion

We observed significant, strong positive correlations between mGluR5 binding and synaptic density in the hippocampus and entorhinal cortex of participants with AD. Cognitively normal participants showed slightly weaker but still strong positive correlations between mGluR5 and synaptic density in the hippocampus only. Whole brain region-based analyses suggested a more widespread pattern of positive correlations between mGluR5 binding and synaptic density due to AD that was not present in older adults with normal cognition. Our findings suggest that loss of mGluR5 in AD may be closely linked to AD related synaptic loss. Further studies may provide insight into the role of mGluR5 at various stages of AD pathologic change, expand our understanding of AD pathogenesis, and aid in the development of novel biomarkers and treatments.

## List of abbreviations

mGluR5: Metabotropic glutamate receptor subtype 5
AD: Alzheimer’s disease
PET: Cognitively normal
CN: Positron emission tomography
*DVR*: Distribution Volume Ratio
*BP*_ND_: Binding Potential Non-displaceable
SV2A: Synaptic vesicle glycoprotein 2A
LTD: Long-term depression
LTP: long-term potentiation
PAM: Positive allosteric modulator
Aβo: Amyloid-β oligomer
PrPc: Cellular prion protein
NIA-AA: National Institute on Aging-Alzheimer’s Association
CDR: Clinical Dementia Rating
MMSE: Mini-Mental Status Examination
aMCI: Amnestic mild cognitive impairment
LM: Logical Memory
SD: Standard deviations
[^11^C]PiB: [^11^C]Pittsburg Compound B
ROI: Region of interest
MRI: Magnetic resonance imaging
MPRAGE: Magnetization-prepared rapid gradient-echo
PVC: Partial volume correction
MOLAR: Motion-compensation OSEM List-mode Algorithm for Resolution-recovery
SRTM2: Simplified reference tissue model–2
*r*: Pearson’s correlation coefficient
FDR: False discovery rate

## Declarations

### Consent for publication

Not applicable

### Availability of data and materials

The data used for these analyses are available from the corresponding author on reasonable request.

### Competing interests

Adam P. Mecca, Richard E. Carson, and Christopher H. van Dyck report grants from the National Institutes of Health for the conduct of the study. Adam P. Mecca reports grants for clinical trials from Eli Lilly and Janssen Pharmaceuticals outside the submitted work. Yiyun Huang reports research grants from UCB and Eli Lilly outside the submitted work. Yiyun Huang, Nabeel B. Nabulsi, and Richard E. Carson have a patent for a newer version of the tracer. Richard E. Carson is a consultant for Rodin Therapeutics and has received research funding from UCB. Richard E. Carson reports having received grants from AstraZeneca, Astellas, Eli Lilly, Pfizer, Taisho, and UCB outside the submitted work. Ryan S. O’Dell reports grants for clinical trials from Cognition Therapeutics and Bristol-Myers Squibb outside of the submitted work. Christopher H. van Dyck reports consulting fees from Kyowa Kirin, Roche, Merck, Eli Lilly, and Janssen and grants for clinical trials from Biogen, Novartis, Eli Lilly, Merck, Eisai, Janssen, Roche, Genentech, Toyama, and Biohaven outside the submitted work.

### Funding

This research was supported by the National Institute on Aging (P30AG066508, P50AG047270, K23AG057794, and R01AG052560, R01AG062276). This publication was made possible by CTSA Grant Number UL1 TR001863 from the National Center for Advancing Translational Science (NCATS), a component of the National Institutes of Health (NIH). The contents of this manuscript are solely the responsibility of the authors, and the funding bodies had no role in the design of the study, data collection, analysis, interpretation, or writing of the manuscript.

### Author contributions

Dr. Salardini and Dr. Mecca had full access to the data and take responsibility for the integrity of the data and the accuracy of the data analysis. Study concept and design: Carson, Huang, Mecca, Nabulsi, O’Dell, Salardini, van Dyck. Acquisition, analysis, or interpretation of data: Carson, Mecca, O’Dell, Salardini, Tchorz, van Dyck. Drafting of the manuscript: Mecca, O’Dell, Salardini, van Dyck. Critical revision of the manuscript for important intellectual content: Carson, Huang, Mecca, Nabulsi, O’Dell, Salardini, van Dyck. Statistical analysis: Mecca, O’Dell, Salardini. Study supervision: Mecca, Salardini, van Dyck.

## Acknowledgements

We thank the research participants for their contributions, and the staff of the Yale ADRU and the Yale PET Center for their excellent technical assistance.

