## Supplemental Materials for "Assessment of the relationship between synaptic density and metabotropic glutamate receptors in early Alzheimer’s disease: a multi-tracer PET study"

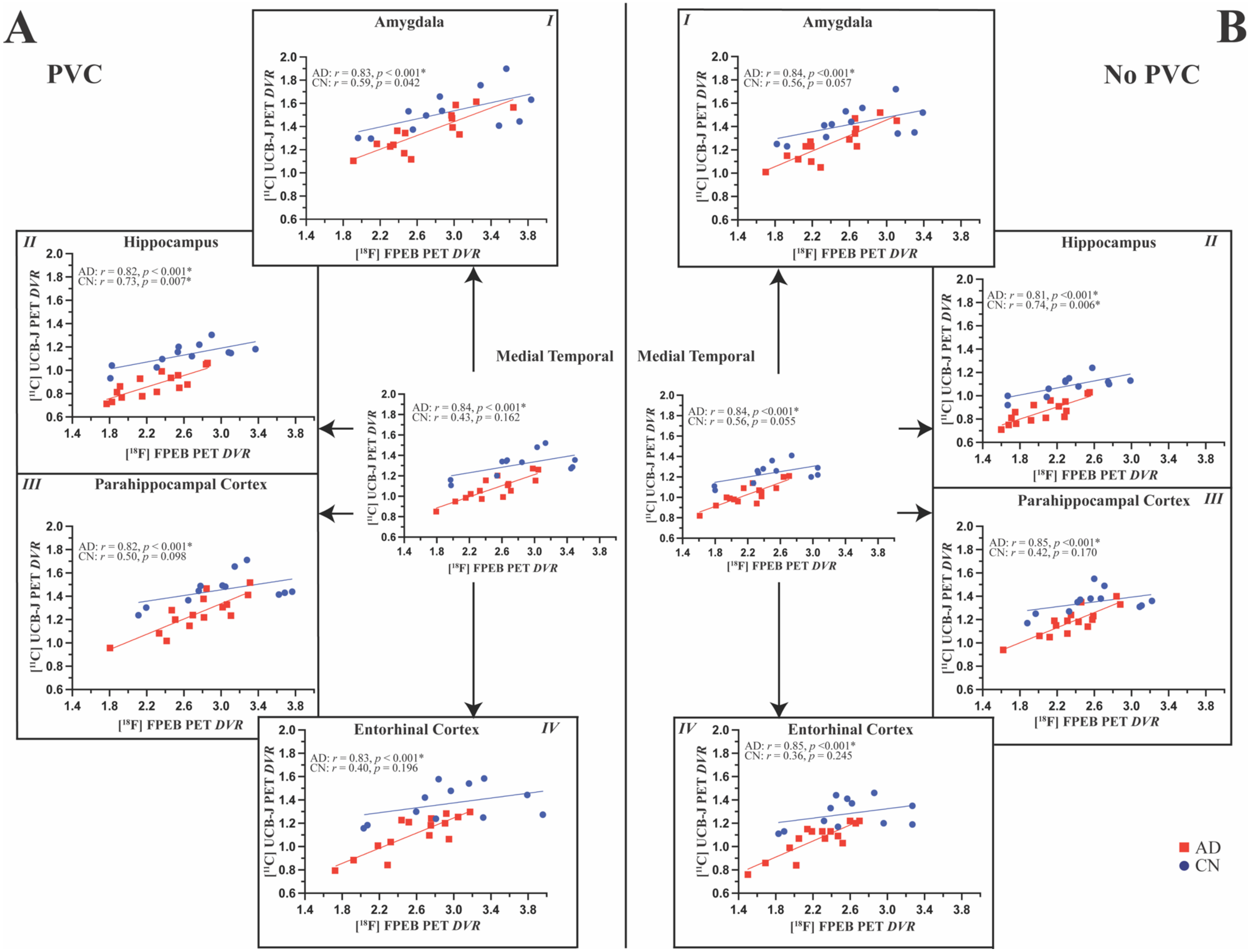


**Supplementary Figure 1. Correlations between mGluR5 and synaptic density in medial temporal regions**

$[^{18}F]$FPEB (mgluR5) and $[^{11}C]$UCB-J (synaptic density) *DVR*s were plotted for each participant. Lines of best fit from univariate regression models are displayed for participants with CN (blue) and AD (red). Pearson's correlation coefficients (*r*) and the associated uncorrected *p* values are displayed for each group. The medial temporal region of interest included bilateral amygdala, hippocampal, parahippocampal cortex, and entorhinal cortex regions. $^{*}p < 0.05$ after FDR correction for multiple comparisons (5 regions per group). Analyses are from 12 participants with normal cognition and 15 participants with AD **(A)** with partial volume correction and **(B)** without partial volume correction of PET data. Abbreviations: AD = Alzheimer's disease; CN = cognitively normal; *DVR* = Distribution volume ratio; FDR = false discovery rate; PVC = partial volume correction; *r* = Pearson's correlation coefficient.

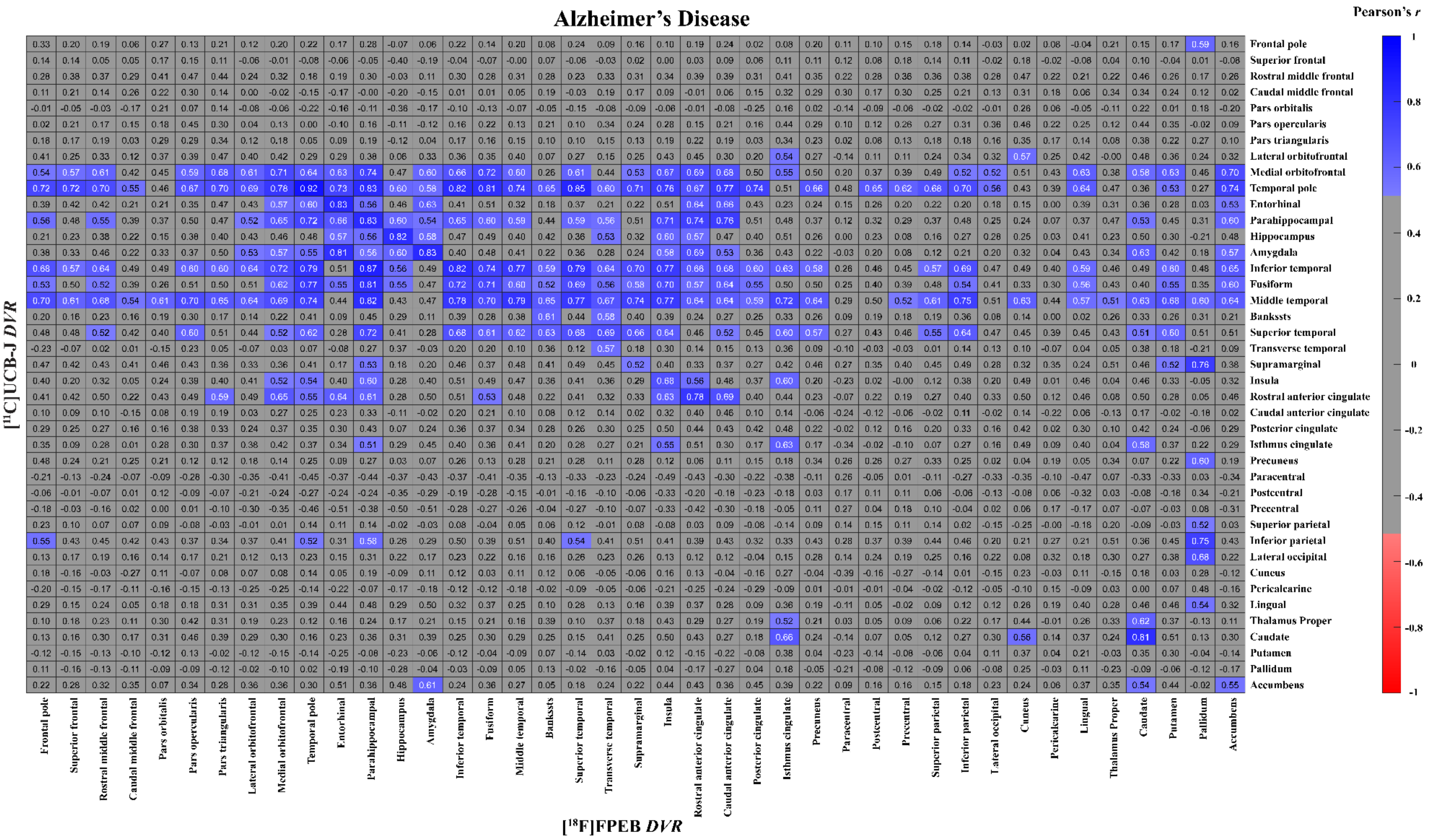


**Supplementary Figure 2. Correlation matrix for mGluR5 and synaptic density (with PVC) of all regions in participants with AD.** The matrix displays Pearson's correlation coefficients ($r$) between [$^{18}$F]FPEB (mGluR5) and [$^{11}$C]UCB-J (synaptic density) PET

*DVRs* for all possible combinations of regions. Data are from 15 participants with Alzheimer's Disease and using PET images that were corrected for partial volume effect. The heat map shows the $r$ for all combinations that had an uncorrected $p < 0.05$.

Abbreviations: AD = Alzheimer's disease; CN = cognitively normal; *DVR* = Distribution volume ratios; mGluR5 = metabotropic glutamate receptor subtype 5; PVC = partial volume corrected

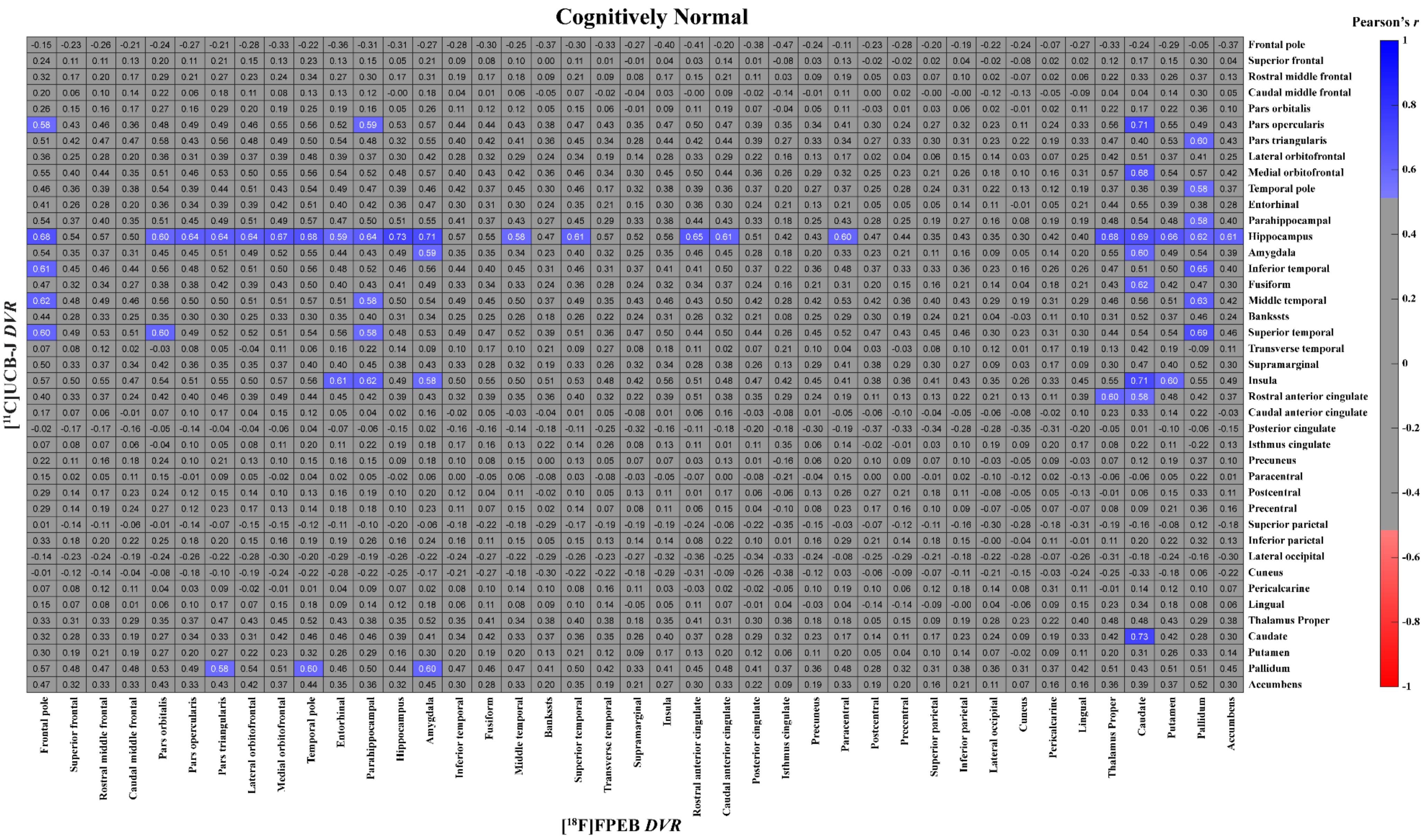


**Supplementary Figure 3. Correlation matrix for mGluR5 and synaptic density (with PVC) of all regions in CN participants.**

The matrix displays Pearson's correlation coefficients ($r$) between [$^{18}$F]FPEB (mGluR5) and [$^{11}$C]UCB-J (synaptic density) PET *DVRs* for all possible combinations of regions.  Data are from 12 cognitively normal participants and using PET images that were corrected for partial volume effect. The heat map shows the *r* for all combinations that had an uncorrected $p < 0.05$. Abbreviations: AD = Alzheimer's disease; CN = cognitively normal; *DVR* = Distribution volume ratios; mGluR5 = metabotropic glutamate receptor subtype 5; PVC = partial volume corrected

### Supplementary Tables and Figures

**Supplementary Table 1. Common AD-affected Regions included in Composite Region of Interest**

| Region | Right Label | Left label |
|---|---|---|
| Hippocampus | 53 | 17 |
| Entorhinal cortex | 2006 | 1006 |
| Parahippocampal cortex | 2016 | 1016 |
| Amygdala | 54 | 18 |
| Rostral anterior cingulate cortex | 2026 | 1026 |
| Caudal anterior cingulate cortex | 2002 | 1002 |
| Posterior cingulate cortex | 2023 | 1023 |
| Isthmus of the cingulum | 2010 | 1010 |
| Precuneus | 2025 | 1025 |
| Frontal pole | 2032 | 1032 |
| Superior frontal gyrus | 2028 | 1028 |
| Rostral middle frontal gyrus | 2027 | 1027 |
| Caudal middle frontal gyrus | 2003 | 1003 |
| Pars orbitals | 2019 | 1019 |
| Pars opercularis | 2018 | 1018 |
| Pars triangularis | 2020 | 1020 |
| Lateral orbitofrontal cortex | 2012 | 1012 |
| Medal orbitofrontal cortex | 2014 | 1014 |
| Inferior temporal gyrus | 2009 | 1009 |
| Temporal pole | 2033 | 1033 |
| Banks of the superior temporal sulcus | 2001 | 1001 |
| Superior temporal gyrus | 2030 | 1030 |
| Transverse temporal gyrus | 2034 | 1034 |
| Fusiform gyrus | 2007 | 1007 |
| Middle temporal gyrus | 2015 | 1015 |
| Superior parietal lobule | 2029 | 1029 |
| Inferior parietal lobule | 2008 | 1008 |
| Supramarginal gyrus | 2031 | 1031 |
| Lateral occipital cortex | 2011 | 1011 |

Individual regions that comprise the composite of regions commonly affected by Alzheimer's disease. FreeSurfer [Version 6.0] Desikan-Killiany atlas label numbers are provided for each hemisphere.

**Supplementary Table 2. Regional correlations between mGluR5 and synaptic density with PVC**

| | Left hemisphere | | | | | | Right hemisphere | | | | | |
|---|---|---|---|---|---|---|---|---|---|---|---|---|
| | **AD** | | **CN** | | **Fisher's *r* to z** | | **AD** | | **CN** | | **Fisher's *r* to z** | |
| **Region** | ***r*** | ***p*** | ***r*** | ***p*** | ***z*** | ***P*** | ***r*** | ***p*** | ***r*** | ***p*** | ***z*** | ***P*** |
| Frontal pole | 0.13 | 0.649 | -0.15 | 0.645 | 0.63 | 0.528 | 0.48 | 0.071 | -0.23 | 0.465 | 1.72 | 0.085 |
| Superior frontal | 0.07 | 0.798 | 0.14 | 0.655 | 0.16 | 0.869 | 0.21 | 0.456 | 0.08 | 0.813 | 0.30 | 0.760 |
| Rostral middle frontal | 0.20 | 0.471 | -0.09 | 0.770 | 0.68 | 0.497 | 0.46 | 0.080 | 0.31 | 0.320 | 0.41 | 0.684 |
| Caudal middle frontal | 0.11 | 0.681 | 0.04 | 0.905 | 0.18 | 0.860 | 0.44 | 0.099 | 0.25 | 0.424 | 0.48 | 0.627 |
| Pars orbitalis | 0.37 | 0.174 | 0.30 | 0.348 | 0.19 | 0.851 | 0.003 | 0.989 | 0.21 | 0.506 | 0.48 | 0.630 |
| Pars opercularis | 0.22 | 0.436 | 0.43 | 0.166 | 0.53 | 0.593 | 0.51 | 0.051 | 0.48 | 0.116 | 0.10 | 0.917 |
| Pars triangularis | 0.24 | 0.382 | 0.49 | 0.101 | 0.67 | 0.502 | 0.45 | 0.089 | 0.52 | 0.083 | 0.20 | 0.842 |
| Lateral orbitofrontal | 0.44 | 0.099 | 0.15 | 0.645 | 0.74 | 0.461 | 0.29 | 0.298 | 0.53 | 0.076 | 0.67 | 0.504 |
| Medial orbitofrontal | 0.74 | 0.002* | 0.50 | 0.093 | 0.87 | 0.383 | 0.66 | 0.007* | 0.54 | 0.072 | 0.44 | 0.656 |
| Temporal pole | 0.94 | <0.001* | 0.35 | 0.258 | 3.09 | 0.002* | 0.88 | <0.001* | 0.60 | 0.040* | 1.57 | 0.115 |
| Entorhinal | 0.82 | <0.001* | 0.51 | 0.087 | 1.34 | 0.181 | 0.81 | <0.001* | 0.20 | 0.541 | 2.08 | 0.037* |
| Parahippocampal | 0.76 | 0.001* | 0.50 | 0.097 | 1.02 | 0.305 | 0.78 | <0.001* | 0.44 | 0.150 | 1.33 | 0.185 |
| Hippocampus | 0.82 | <0.001* | 0.68 | 0.014 | 0.77 | 0.441 | 0.79 | <0.001* | 0.78 | 0.002* | 0.05 | 0.958 |
| Amygdala | 0.69 | 0.005* | 0.51 | 0.093 | 0.64 | 0.519 | 0.88 | <0.001* | 0.61 | 0.036* | 1.57 | 0.117 |
| Inferior temporal | 0.70 | 0.004* | 0.45 | 0.144 | 0.86 | 0.387 | 0.72 | 0.002* | 0.43 | 0.161 | 1.02 | 0.309 |
| Fusiform | 0.54 | 0.037* | 0.29 | 0.354 | 0.69 | 0.491 | 0.79 | <0.001* | 0.34 | 0.283 | 1.62 | 0.105 |
| Middle temporal | 0.66 | 0.007* | 0.39 | 0.207 | 0.87 | 0.383 | 0.75 | 0.001* | 0.56 | 0.056 | 0.75 | 0.452 |
| Bankssts | 0.53 | 0.043* | -0.38 | 0.222 | 2.24 | 0.025* | 0.55 | 0.033* | 0.48 | 0.116 | 0.23 | 0.820 |
| Superior temporal | 0.58 | 0.024* | 0.40 | 0.201 | 0.55 | 0.585 | 0.69 | 0.004* | 0.61 | 0.034* | 0.32 | 0.748 |
| Transverse temporal | 0.33 | 0.230 | 0.12 | 0.714 | 0.51 | 0.612 | 0.74 | 0.002* | 0.51 | 0.089 | 0.87 | 0.383 |
| Supramarginal | 0.46 | 0.082 | 0.20 | 0.527 | 0.67 | 0.504 | 0.49 | 0.062 | 0.35 | 0.260 | 0.39 | 0.699 |
| Insula | 0.52 | 0.047* | 0.32 | 0.314 | 0.56 | 0.575 | 0.74 | 0.002* | 0.69 | 0.012* | 0.21 | 0.832 |
| RAC | 0.85 | <0.001* | 0.50 | 0.100 | 1.63 | 0.103 | 0.68 | 0.005* | 0.33 | 0.296 | 1.13 | 0.259 |
| CAC | 0.54 | 0.038* | 0.28 | 0.371 | 0.71 | 0.480 | 0.36 | 0.182 | 0.08 | 0.811 | 0.69 | 0.490 |
| Posterior cingulate | 0.27 | 0.321 | -0.21 | 0.505 | 1.13 | 0.257 | 0.51 | 0.052 | -0.11 | 0.730 | 1.53 | 0.126 |
| Isthmus cingulate | 0.38 | 0.163 | 0.48 | 0.113 | 0.28 | 0.775 | 0.77 | 0.001* | 0.24 | 0.455 | 1.79 | 0.074 |
| Precuneus | 0.26 | 0.346 | -0.04 | 0.885 | 0.71 | 0.475 | 0.48 | 0.071 | 0.15 | 0.638 | 0.83 | 0.404 |
| Paracentral | 0.17 | 0.544 | 0.30 | 0.345 | 0.31 | 0.757 | 0.39 | 0.150 | 0.03 | 0.914 | 0.86 | 0.392 |
| Postcentral | 0.08 | 0.768 | 0.30 | 0.344 | 0.51 | 0.609 | 0.27 | 0.330 | 0.24 | 0.456 | 0.08 | 0.938 |
| Precentral | 0.18 | 0.518 | 0.17 | 0.591 | 0.02 | 0.984 | 0.21 | 0.444 | 0.16 | 0.616 | 0.12 | 0.902 |
| Superior parietal | 0.15 | 0.581 | -0.17 | 0.602 | 0.74 | 0.460 | 0.13 | 0.651 | -0.009 | 0.975 | 0.31 | 0.755 |
| Inferior parietal | 0.00 | 0.980 | 0.10 | 0.749 | 0.22 | 0.826 | 0.62 | 0.013* | 0.19 | 0.548 | 1.21 | 0.227 |
| Lateral occipital | 0.12 | 0.677 | -0.21 | 0.505 | 0.76 | 0.448 | 0.31 | 0.253 | -0.21 | 0.506 | 1.23 | 0.219 |
| Cuneus | 0.25 | 0.364 | -0.16 | 0.606 | 0.96 | 0.335 | 0.31 | 0.259 | 0.02 | 0.947 | 0.68 | 0.496 |
| Pericalcarine | 0.27 | 0.338 | 0.42 | 0.172 | 0.40 | 0.688 | 0.41 | 0.125 | 0.16 | 0.623 | 0.64 | 0.524 |
| Lingual | 0.33 | 0.227 | 0.06 | 0.855 | 0.65 | 0.517 | 0.48 | 0.070 | 0.27 | 0.391 | 0.55 | 0.582 |
| Thalamus | 0.30 | 0.283 | 0.24 | 0.455 | 0.14 | 0.887 | 0.38 | 0.164 | 0.63 | 0.026* | 0.80 | 0.424 |
| Caudate | 0.80 | <0.001* | 0.70 | 0.011* | 0.55 | 0.580 | 0.74 | 0.001* | 0.70 | 0.011* | 0.20 | 0.842 |
| Putamen | 0.35 | 0.194 | 0.26 | 0.406 | 0.23 | 0.820 | 0.19 | 0.486 | 0.24 | 0.452 | 0.11 | 0.914 |
| Pallidum | 0.22 | 0.434 | 0.59 | 0.041* | 1.05 | 0.293 | 0.03 | 0.926 | 0.38 | 0.222 | 0.85 | 0.396 |
| Accumbens area | 0.43 | 0.108 | 0.03 | 0.919 | 0.97 | 0.331 | 0.45 | 0.091 | 0.58 | 0.045* | 0.42 | 0.676 |
| Ventral Diencephalon | 0.27 | 0.326 | 0.06 | 0.855 | 0.50 | 0.617 | 0.53 | 0.040* | 0.10 | 0.752 | 1.12 | 0.263 |

Pearson's *r* and associated *p* value is reported for the correlation between partial volume corrected [$^{18}$F]FPEB PET (mGluR5) and [$^{11}$C]UCB-J PET (synaptic density) *DVR* in each brain region. Fisher *r*-to-*z* transformation was used to compare correlation coefficients of CN and AD groups. The data were from 12 CN participants 15 participants with AD. * $p < 0.05$. Abbreviations: AD = Alzheimer's disease; CN = cognitively normal; CAC = Caudal Anterior Cingulate; *DVR* = Distribution volume ratio; PVC = Partial volume correction; RAC = Rostral Anterior Cingulate
